# *O*-GlcNAc transferase contributes to sex-specific placental deregulation in gestational diabetes

**DOI:** 10.1101/2022.05.24.492568

**Authors:** Yiwen Cui, Meredith Cruz, Anna Palatnik, Stephanie Olivier-Van Stichelen

**Affiliations:** Department of Obstetrics and Gynecology, Medical College of Wisconsin, Milwaukee, WI, 53226; Cardiovascular Center, Medical College of Wisconsin, Milwaukee, WI, 53226; Department of Biochemistry, Medical College of Wisconsin, Milwaukee, WI, 53226

**Keywords:** *O*-GlcNAcylation, Placenta, RNA-sequencing, gestational diabetes

## Abstract

Fetal sex has been known to be associated with differential risks for adverse perinatal outcomes. Recently it has been highlighted as a risk factor for gestational diabetes (GDM), which carries short- and long-term cardiometabolic implications for the pregnant women and their offspring. Understanding the molecular mechanisms through which fetal sex can modulate placenta physiology can help identify novel molecular targets for future clinical applications. In this study, we investigated the nutrient-sensing *O*-GlcNAc pathway as a potential mediator of sex-driven placenta dysfunction due to the unique location of the *O*-GlcNAc transferase (OGT) enzyme on the X chromosome. Using banked placenta tissues, we demonstrated that placenta *OGT* is downregulated in women with GDM, especially when carrying a male offspring. We modeled our observation *in vitro* using male placenta trophoblast BeWo cells in which OGT was knocked down. A comprehensive transcriptomic profile *via* RNA sequencing demonstrated changes in hormonal, inflammatory and immunologic markers, toward GDM-like transcriptional signatures. Altogether, this study suggests that OGT deserves important consideration for sexual dimorphism observed in GDM and highlights the importance of O-GlcNAcylation in placenta endocrine physiology.

## INTRODUCTION

The placenta is a fetal secretory organ responsible for producing hormones, cytokines, and regulatory proteins for the adaptation of maternal physiology necessary for pregnancy. Aberrant placenta function leads to various endocrine diseases, the most common being gestational diabetes (GDM) estimated to affect one in ten pregnancies globally^1^. The diagnosis of GDM poses immediate risks to the pregnancy, including high likelihood for preeclampsia (preE), stillbirth and preterm delivery. Furthermore, GDM also carries long-term health risks for the mother and offspring^2–5^. Up to 70% of women with a history of GDM will develop type 2 diabetes (T2DM) within 10 years following pregnancy ^6^, while the offspring is at increased risk for fetal adiposity, obesity and early-onset T2DM as young adults^7–11^.

Recent clinical evidence suggested that fetal sex is an additional risk factor for GDM, as women who carry a male fetus have up to a 39% odds risk for developing GDM (12). Conversely, those who develop GDM while carrying a female fetus may have the higher inherent risk toward insulin resistance ^12,13^. Thus, fetal sex has also been proposed as a predictor for a woman’s risk for postpartum progression from GDM to T2DM. This sexual dimorphic risk for GDM provides a unique comparative framework to study inherent metabolic drivers for glucose and insulin deregulation and, by extension, identify targets for metabolic therapy.

In this paper, we propose that the fetal placental *O*-GlcNAcylation pathway widely disrupts placenta function leading to a sexually dimorphic risk for the development of GDM. *O*-GlcNAcylation is a highly conserved modification consisting of the post-translational addition of a single residual of N-acetylglucosamine onto serine or threonine moieties of intracellular proteins (Figure 1A)^14^. This modification is highly dependent on glucose level and consequently, the pathway is considered an important nutrient sensor, capable of regulating a wide range of physiologic processes in response to nutritional environment ^15–17^.

**Figure 1:**
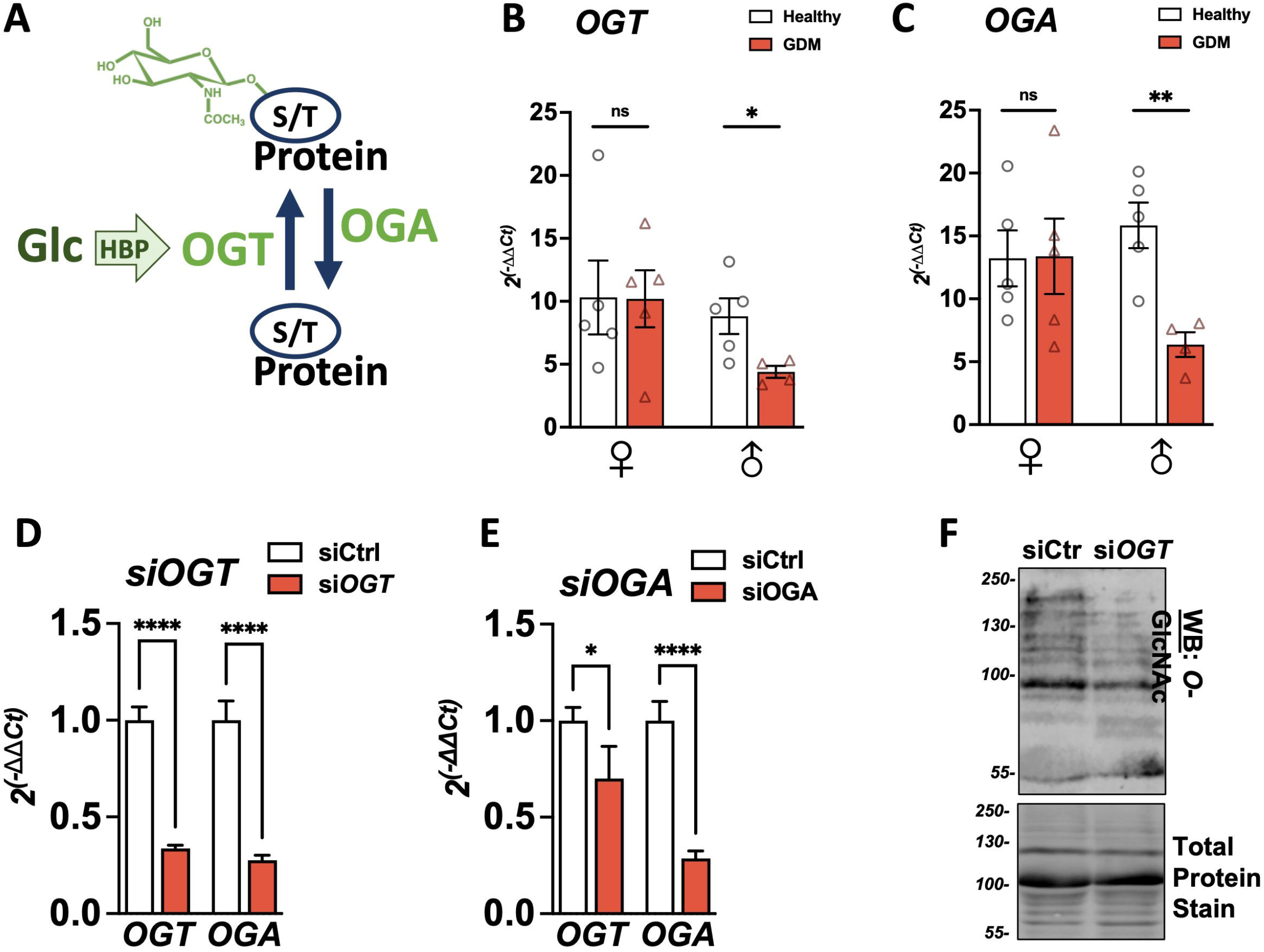
GDM correlates with downregulation of O-GlcNAc enzymes in male placentas. *O*-GlcNAcylation of proteins. **A** 2-3% of glucose (Glc) entering cells is shuttled through the Hexosamine Biosynthesis Pathway (HBP) to produce UDP-GlcNAc, the nucleotide donor for *O*-GlcNAcylation. *O*-GlcNAc transferase (OGT) adds the GlcNAc residue to serine or threonine (S/T) residues. *O*-GlcNAcase (OGA) removes the sugar residue. **B/C** Expressions of O-GlcNAc transferase (OGT) (B) or O-GlcNAcase (OGA) (C) were measured in healthy or Gestational diabetes (GDM) placentas by qPCR and reported on ACTIN. **D/E** *OGT* and *OGA* expression were measured by qPCR in BeWo cells transfected with siRNA against *OGT* (D) or *OGA* (E) for 48h. Relative expression was calculated using the -2ΔΔCt method. Significance was calculated by student t-test; ns ≥ 0.05, * p < 0.05, ** p < 0.01, *** p < 0.001, **** p < 0.0001. **F** *O*-GlcNAcylation levels in BeWo cells transfected with siRNA against *OGT* for 48h. Total protein stain was used as the loading control.

*O*-GlcNAcylation has already been associated with X-linked diseases^18–22^ due to the localization of the main enzyme *O*-GlcNAc transferase (OGT) on the X-chromosome (Xq13.1)^23,24^. Numerous studies have found *Ogt* to escape X-chromosome inactivation, potentially resulting in differential protein dosages ^25–28^. The extra copy of Ogt can lead to increased plasticity against environmental stress factors, as shown in pregnant murine models (30). Particularly, during environmental stress from hyperglycemia, the extra copy of Ogt has been shown to offer increased plasticity in response in pregnant murine models (30). Finally, animal phenotypes resulting from *O*-GlcNAc modulation varied based on sex^29–36^, emphasizing the link between sexual dimorphism and *O*-GlcNAcylation.

Herein, we investigated the role of OGT in the development of GDM. We first demonstrated a male-specific decrease in *OGT* and *OGA* expression from human placenta samples collected from women with GDM when compared to healthy controls. We next modeled the decreased *OGT* expression *in vitro* and performed a comprehensive transcriptomic profile using RNA sequencing. We identified significant changes in the production of placenta hormones and inflammatory markers implicated in insulin dysregulation of GDM. Altogether, this study demonstrates a role for OGT and O-GlcNAcylation in placenta endocrine deregulation and might explain the sexual dimorphism observed in GDM outcomes.

## RESULTS

### OGT is selectively downregulated in male fetal placenta from GDM mothers

To examine the pathological relevance of *O*-GlcNAcylation in GDM based on fetal sex, *OGT* and *OGA* expression were measured by qPCR in human placentas grouped by fetal sex and maternal diagnosis of GDM (n=4+/group) (Table 1). *OGT* and *OGA* were significantly downregulated in male GDM placenta compared to control (*OGT*: p=0.03, *OGA* p=0.004, Figure 1 B/C). In contrast, this difference was not observed in the female group.

There were significant differences in maternal age and body mass index (BMI) between GDM and control subgroups within each sex (Table 1). However, OGT and OGA levels poorly correlate with maternal characteristics (Figure S1), suggesting that the significant decrease in *OGT* observed in the male group is related to GDM pathology.

To determine the physiologic impact of *O*-GlcNAc enzymes downregulation toward GDM development, we established a cell culture model using BeWo cells, which are female choriocarcinoma placenta cells sharing the secretory profile of placenta syncytiotrophoblasts. We knocked down OGT and OGA individually using small interfering RNA (siRNA) (Figure 1 D/E); control cells were treated with a Negative control scramble siRNA (SiCtrl). While both siRNAs were successful, *OGT* knocked down led to a more substantial decrease in *OGA* level, closer to the levels observed in Healthy *vs*. GDM placentas. Furthermore, si*OGT* led to a strong reduction in the total *O*-GlcNAcylation level observed by Western Blot (Figure 1 F). This confirmed literature reports that OGA expression can fluctuate with OGT modulation (38). We subsequently focused on the siOGT model as it closely simulated our findings from the placenta samples.

### Modulating OGT leads to critical changes in the placental transcriptome

Using the previously established cellular model, six RNA-Seq datasets were generated from two sample groups (siCtrl, siOGT) to conduct comparative pathway analysis that could elucidate the role of OGT downregulation in placental dysfunction leading to GDM. Each sample group was provided in biological replicates and obtained a total of 298,978,153 reads (44-55 million reads per sample) with ∼92% were over 30 bases (Table S1). Particularly, 68 genes (48 upregulated and 20 downregulated) had more than a 2-fold change with an adjusted p-value (AdjpValue) < 0.05 (Figure 2A, Table S2). An additional 4,383 genes were also significantly deregulated (AdjpValue <0.05) but with less than a 2-fold change (Table S2).

**Figure 2:**
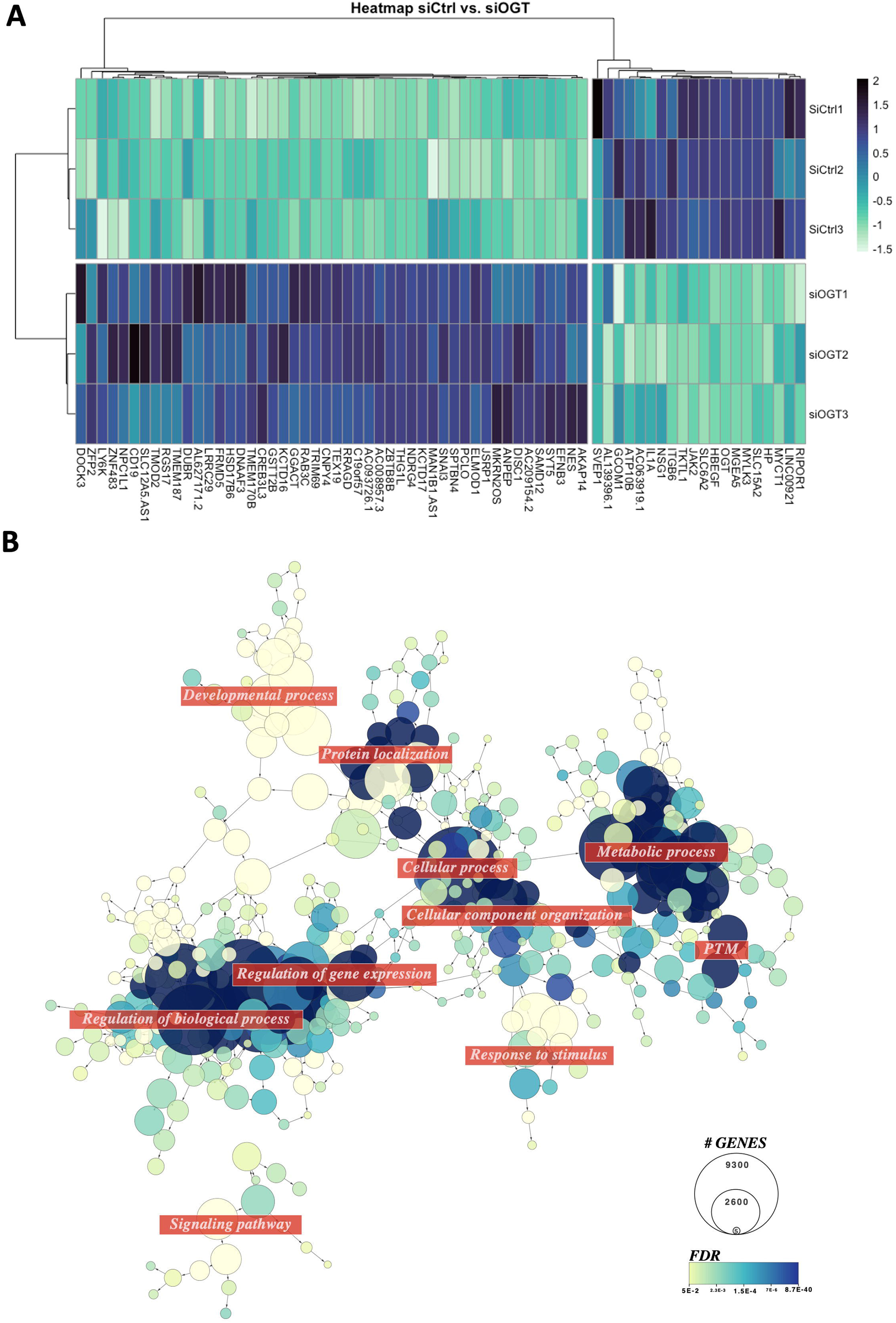
Modulating OGT in male placental cells leads to important changes in the transcriptome. **A** Heatmap of the deregulated genes in siOGT vs. siCtrl Bewo cells. Genes shown have an adjPValue<0.05 and abs(Log2FC)>1. **B** Network representation of enriched GO terms (FDR<0.05) using deregulated genes (AdjPValue<0.05) in siOGT vs. siCtrl Bewo Cells.

Gene Ontology (GO) analysis on the significatively deregulated genes demonstrated enrichment in the cellular metabolic process (GO:0044237, FDR=8.78E-40), comprising protein post-translational modification (GO:0043687, FDR=2.12E-10) and gene expression (GO:00010467, FDR=1.83E-14), and subsequent regulation of cellular process (GO:0050794, FDR=3.91E-11), encompassing, regulation of gene expression (Figure 2B, Table S3). Organelle organization (GO:0006996, FDR=1.92E-22) was also enriched driven by chromatin modification (GO:0016568, FDR= 1.46E-09). Macromolecule localization (GO:0033036, FDR=5.58E-13), including protein transport (GO:0015031, FDR=9.35E-11) and cellular localization (GO: 0051641, FDR= 3.27E-09) was also associated with the significantly deregulated genes in si*OGT* cells. Overall, this analysis demonstrated the validity of our model as it highlighted the previously described role of *O*-GlcNAcylation in transcriptional regulation^14^.

### Gene expression profile from siOGT BeWo cell line overlaps with transcriptomic changes observed in GDM placenta

We further compared the previous RNA-sequencing data with previously published transcriptomes of human GDM placenta. Inputting the deregulated genes in si*OGT* (AdjPValue<0.05) to the Alliance of Genome Resources database (https://www.alliancegenome.org) and querying for GDM-related genes (Disease Ontology (DO)ID:111714), we found that our model shared 32% of gene expression (10/ 31 genes) (Figure 3A). Interestingly, the siOGT model also overlapped in 21% of preE gene deregulations, highlighting potentially shared pathways between preE and GDM and the involvement of OGT in both pathologies for future consideration (Figure S2).

**Figure 3:**
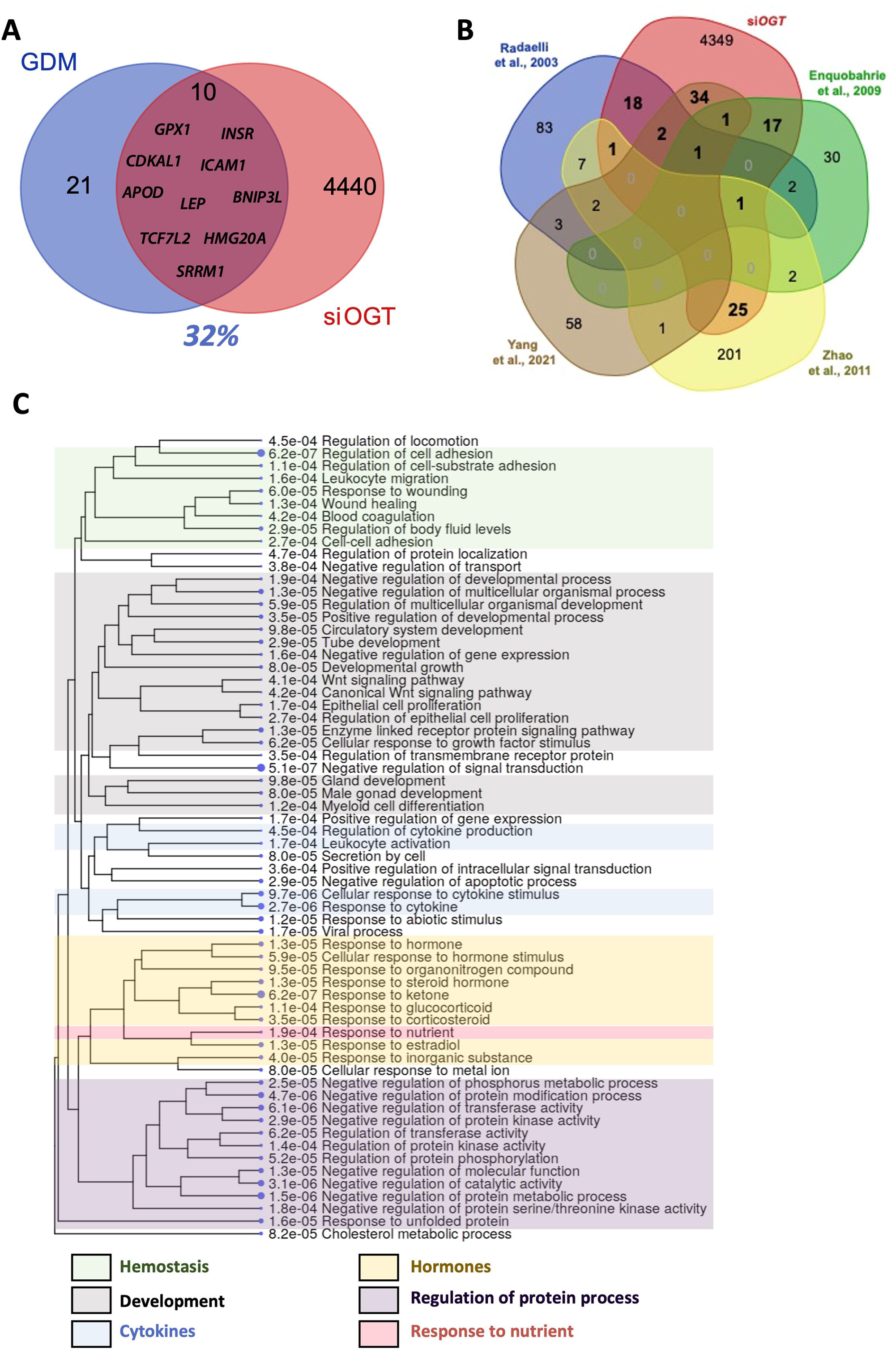
*OGT* siRNA in BeWo cells lead to transcriptional changes common with GDM placentas. **A** GDM signature genes were found in the Alliance of Genome Resources under DOID: 11714. **B** Venn diagram representing commonly deregulated genes in placenta from reference GDM studies. **C** GO enrichment tree of the 100 deregulated genes found in our study and previously identified to be deregulated in GDM placentas.

As the Alliance of Genome Resources database does not offer tissue-specificity for the GDM-related gene, we cross-referenced si*OGT* deregulated genes with reference studies specifically reporting GDM transcriptional changes in human placenta ^37–39^ or purified trophoblasts ^40^. While neither *OGT* nor *OGA* was significantly deregulated in these studies, none of these studies reported nor stratified their analysis by fetal sex. Nevertheless, 100 shared deregulated genes (Figure 3B, Table S4) were commonly found in these studies and ours. Go enrichment of the shared genes highlighted the following categories: hemostasis, cytokine signaling, peptide and steroid hormone signaling, developmental processes, and response to nutrients (Figure 3C).

For each GO term identified previously, a volcano plot of transcriptional changes in siOGT vs. siCtrl was plotted to investigate the importance of OGT in each pathway (Figure 4). Like T2DM, GDM is a complex integration of hormonal, immune, and inflammatory mechanisms, all deregulated in si*OGT* BeWo cells (Figure 4). More specifically, one of the fundamental functions of the placenta is to produce hormones that drive metabolic adaptations in pregnancy. We found that downregulation of *OGT* directly impacted the production of several essential hormones, leaning toward a diabetogenic profile (Figure 4F). Consequently, we independently confirmed that GH2, hPL, hCG, and LEP were similarly deregulated by siRNA OGT in BeWo Cells (Figure S3).

**Figure 4:**
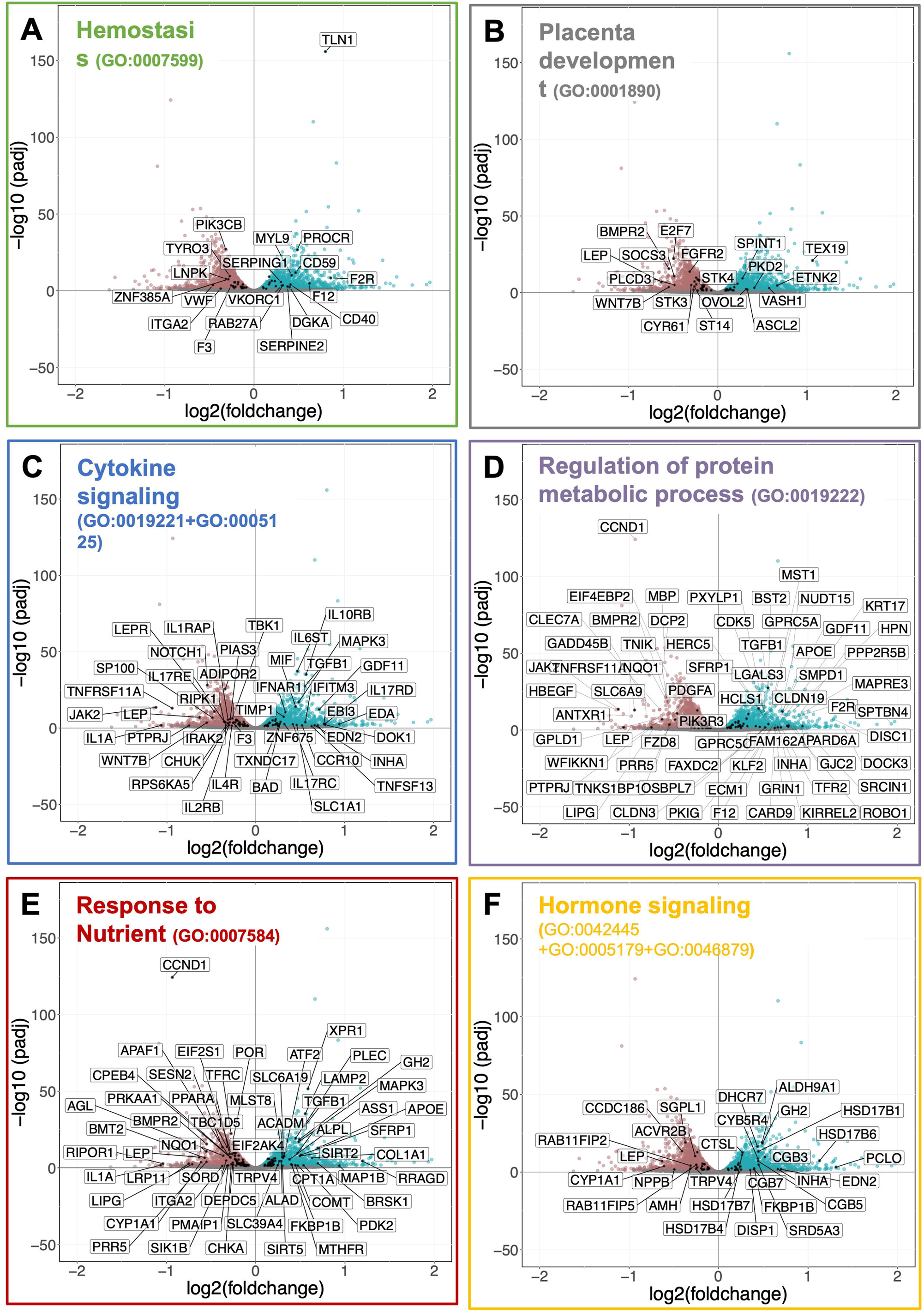
Various pathways are deregulated in siOGT BeWo cells compared to Ctrl, including nutrient sensing and hormone signaling. Volcano plots for siCtrl vs. si*OGT*. Colored datapoints have an AdjpValue<0.05. Genes are labeled when they are significant and found in related GO terms.

### Transcriptomic changes in siOGT cells can be explained by common transcription factors

We further constructed a regulatory network from the deregulated genes from the siRNA, which can offer potential clinical application as biomarker or for therapy. We interrogated the TRRUST database with the list of significantly deregulated genes in si*OGT* (AdjpValue<0.05) associated with hormone-related GO terms (GO:0042445, GO:0005179, GO:0046879) and discovered that transcription factors such as *SP1, STAT1*, and *NFKB1* were responsible for the transcription of a large portion of deregulated genes and were also themselves significantly deregulated in si*OGT* BeWo cells compared to siCtrl (Figure 5A/B). We also found other transcription factors like *CEBPA/B, REST, CREB1, RELA*, TFAP2A, and *FOXO1* that were responsible for regulating the transcription of many deregulated genes in si*OGT* but were themselves not significantly altered by *OGT* knockdown (Figure 5A/B). All but CEBPA were identified as *O*-GlcNAcylated by the *O*-GlcNAc (Figure 5A) while most were expressed in placentas, with the most specificity for *TFAP2A* (Figure 5B). Finally, *NR5A1, JUN*, and *GATA4* were poorly represented in placentas despite their association with a small percentage of deregulated genes, suggesting that they are unlikely to be relevant to placental deregulation toward GDM (Figure 5A/B).

**Figure 5:**
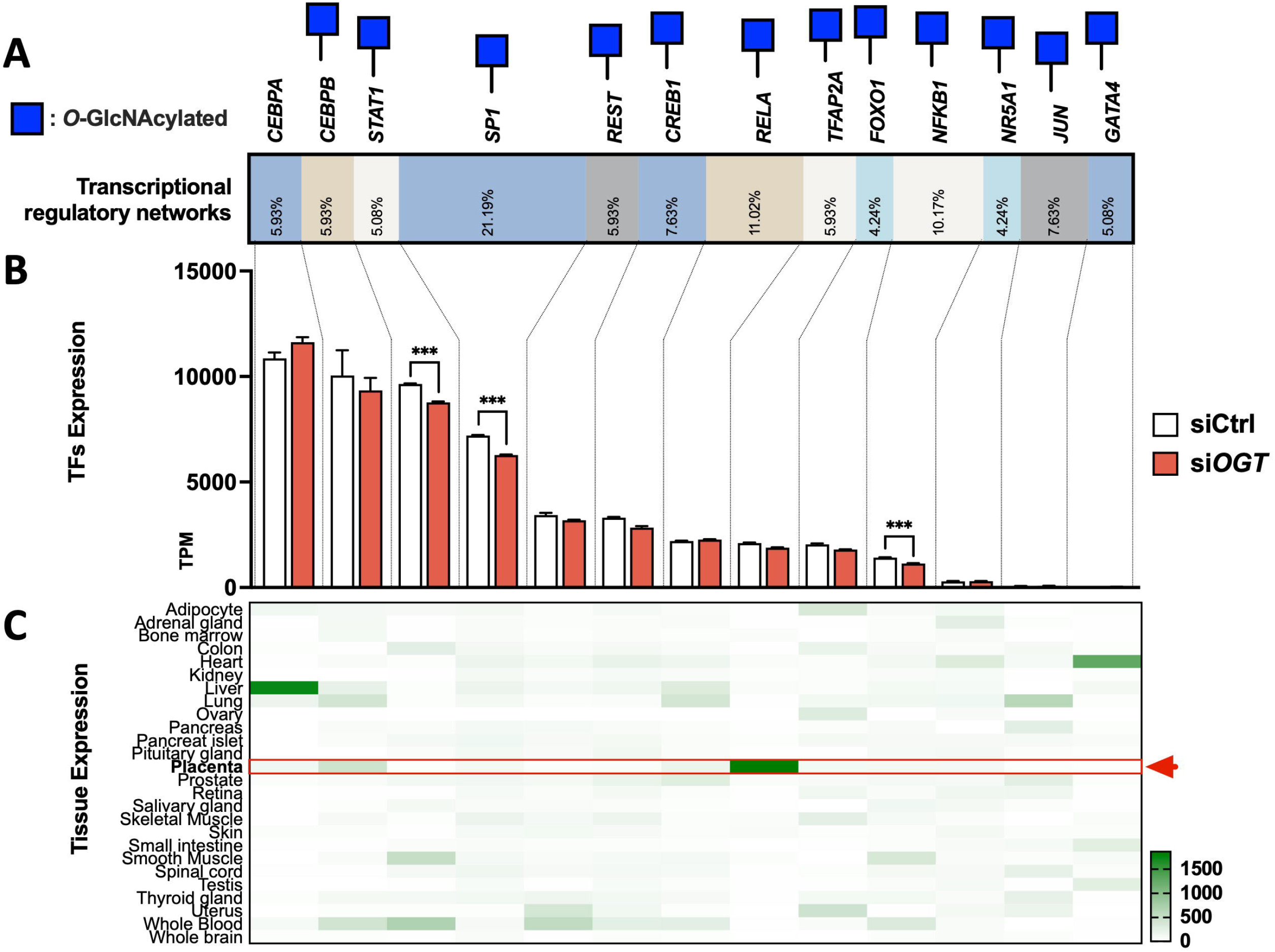
Common transcription factors explained hormonal changes in expression levels after si*OGT* in BeWo cells. **A** TRRUST analysis highlighted the transcription factor responsible for regulating hormones-related genes deregulated in siOGT. Blue Square represents the *O*-GlcNAcylation status assessed with the *O*-GlcNAcylated database. **B** RNA-sequencing expression of transcription factors identified by the TRRUST database. TPMs are represented for biological replicate, and significance was analyzed by regular T-Test. *** p < 0.001. **C** Tissue expression for each transcription factor was assessed in BioGPS and normalized by transcription factors.

## DISCUSSION

Sex differences exist throughout life, particularly when gestational exposures can impact disease susceptibility^41^. Pregnancies with male and female fetuses have differing obstetric risk profiles including miscarriage, preterm birth, and placenta insufficiency^42–44^. As the placenta mediates fetal growth and underlies many pregnancy complications, sex differences arising *in utero* can influence placenta development and function.

This study shows how a sex-chromosome gene can have broad impacts on placenta trophoblasts to activate hormonal, endocrine, and immunologic pathways diving toward insulin resistance and likely, maternal gestational diabetes. Our findings corroborate the clinical data supporting the risk of carrying a male fetus for gestational diabetes.

### O-GlcNAcylation is deregulated in GDM placentas of male origins

Our study focused on the X-linked gene *OGT* in the O-GlcNAcylation pathway, which has been suggested in the literature to escape X-inactivation in the placenta and, consequently, result in higher levels of the enzyme in female versus male placentas^25–28^. In studies on murine prenatal stress, males’ placental OGT and *O*-GlcNAcylation levels were more impacted than females. With extra copies of *OGT* available, it was hypothesized that female offspring may have a greater capacity to rapidly respond to maternal stress^29^ or, in the case of gestational diabetes, prolonged exposure to hyperglycemic stress^45–48^. In this context, we hypothesized that male placental *OGT* and *O*-GlcNAcylation levels might be more affected by stressors leading to induction of GDM. Using matched human placental tissue of male or female origins, we show that, as expected, *OGT* and also *OGA* were downregulated in male GDM placentas compared to matched healthy samples but not in female offspring placenta. This suggests that males and females will demonstrate variable responses to *O*-GlcNAc-dependent processes. Because *O*-GlcNAc signaling is involved in many physiological and pathological processes, our novel observation emphasizes a potential responsibility for *O*-GlcNAcylation in observed sexual dimorphisms in GDM outcomes ^12,13,49^.

Previous studies have investigated the role of *O*-GlcNAcylation in the placenta, using deletion models of *Ogt* in trophoblast lineage in murine models. Consequences of placental-specific *Ogt* deletion on metabolic homeostasis included increased responsivity of the hypothalamic-pituitary-adrenal stress axis in male offspring and increased insulin resistance to high-fat-diet in females ^36,50^. However, while rodent models have enabled significant advances in understanding the role of *O*-GlcNAcylation in placenta function, it is an imperfect simulation of human placental biology and pathophysiology. Indeed, mice placenta has a much more limited endocrine role; it secretes a significantly lower level of steroids, including a complete absence of GH2 and continuous secretion from original endocrine tissues throughout pregnancy, *e*.*g*., corpus luteum (progesterone) or the pituitary gland (GH)^51^. This lack of endocrine function in mice placentas accentuate the role of hormones in pregnancy complications like GDM as the mice never spontaneously develop GDM^52^. However, mice supplemented with human placenta hormones (hGH^53,54^ and hPL^55,56^, steroid hormones^57–62^) develop similar insulin resistance as human.

Thus, human-based models are needed to accurately recapitulate the placenta endocrine role in disease, hence the use of human syncytiotrophoblasts in the current study.

### OGT downregulation drives GDM-like transcriptomic changes

To demonstrate the link between *OGT* downregulation seen in male GDM placenta and GDM development, we modeled our finding in an engineered BeWo cell culture. Comparing the resultant gene changes, we found an over 30% overlap gene expression with previously published GDM-associated genes. Notable among the shared genes are the transcription factor 7-like 2(TCF7L2) and cdk5 regulatory associated protein 1-like 1 (CDKAL1), two of the most potent genes associated with T2DM consistently replicated in multiple populations and implicated in impaired glucose-mediated insulin secretion^63,64^.

Interestingly, cross-referencing the deregulated genes in si*OGT* BeWo with other transcriptomic profiling of GDM placentas^37–40^ revealed disturbance in placental hormone signaling, previously stressed as one of the causes of GDM. Indeed, while proper placental hormones secretion is necessary to reach a natural state of mild insulin resistance in pregnancy to meet fetal nutritional needs^65–79^, dysregulation of hormonal secretion, particularly placental growth hormones (GH2)^53,54,66^, Lactogen (PL)^55,56^, estrogens and progesterone^80–82^, and cytokines ^37,83^, are thought to influence the development of GDM. It is the placenta production of such hormones^53–56,66,71,80–82,84–86 58,79,87^, unique to pregnancy, that has been suggested to contribute to the key clinical characteristic of transiency in GDM, vastly different from the slow progression in T2DM.

Following si*OGT*, we noted an increase in placenta GH, which is a potent insulin antagonist that stimulates maternal lipolysis and hepatic gluconeogenesis^88^. Its rise in the second half of gestation, corresponding to the emergence of insulin resistance during pregnancy, is thought to participate in GDM etiology. Physiologically, this rise in insulin resistance is countered by the upregulation of maternal insulin production driven by placenta-produced prolactin (hPL). The surge in hPL parallels an increase in beta-cell replication in pregnancy, which has been attributed to the role of hPL in prolonging beta-cell survival in isolated human and rodent islets^89,90^. However, si*OGT* cells did not compensate the increased GH2 production with increased hPL. Additionally, the significant decrease in leptin and leptin receptors in the si*OGT* cells further contributes toward a decompensatory capacity to respond to GH-driven insulin resistance. Leptin is consistently shown to be insulin-sensitizing ^91,92^ and its rise in pregnancy is driven by placental trophoblasts ^93^. Thus, we demonstrate that downregulation of *OGT* may contribute to insulin resistance as a consequence of GH hyperproduction in placenta trophoblasts, while limiting compensatory insulin sensitivity measures (hPL and Leptin), altogether leaning toward a hormonal diabetogenic profile.

Beyond the role of placenta-produced hormones, the development of gestational diabetes is also multifactorial with inflammatory, vascular and immunologic factors. This is evident in the shared gene expression between GDM and preE ^94^, also reflected in our si*OGT* model. Thus, it is not surprising that a role for *O*-GlcNAc signaling has been proposed in preeclampsia as well ^95^. We observed the upregulation of inflammatory genes for IL-6, -10, and -17 for an overall skew toward inflammation, as evidenced by the increased Macrophage Migration Inhibitory Factor (MIF) expression in si*OGT* BeWo cells. MIF is recognized as one of the first molecules to arrive at an inflammation site and is critical in proinflammatory innate immune responses. MIF has been implicated in immunological and inflammatory diseases such as rheumatoid arthritis^96^, obesity^97,98^, and diabetes^99,100^. As a center of MIF production, the placenta can directly impact pancreatic β-cell viability and functionality. Under inflammatory conditions, MIF can lead to β cell apoptosis^101^, while the downstream inflammatory signaling can further create a toxic microenvironment hindering the physiologic expansion of β cell mass expected in pregnancy^102,103^.

### Placental endocrine deregulation in GDM might be due to the O-GlcNAcylation of specific transcription factors

In our constructed regulatory network of hormonal genes in trophoblast cells, we identified several critical O-GlcNAc-regulated transcriptional factors (TFs) for future focus. While some of them are widely studied and ubiquitous TFs like SP1^104^, TFAP2A is a highly placental-specific transcription factor responsible for 6% of hormonal gene regulations. Its role in regulating the expression of placental endocrine genes, including *hCG*^105^, CYP11A1^106(p1)^, and HSD17s^107^, is particularly relevant as they are altered by siOGT. Interestingly, abnormal TFAP2A protein levels have been found in pathologic placentas, including preE, GDM, chronic hypertension, and Fetal Growth Restriction(FGR)^108^. Furthermore, via the *O*-GlcNAc database, we were able to identify one study in which TFAP2A was found to be *O*-GlcNAcylated in breast cancer^109(p1),110^. The deregulation of TFAP2A post-transcriptionally modulated by *O*-GlcNAcylation can lead to broad downstream placental hormonal changes characteristic of GDM.

In summary, our study provides new insights into how *O*-GlcNAc-dependent sexual dimorphism can drive placental pathways toward pregnancy complications like GDM. Understanding the biological mechanisms underlying the sex differences in pregnancy outcomes becomes important as gestational diseases like preE and GDM, previously thought to be transient, have lasting effects on the health of offspring and mother decades after the event. As clinical practice evolves towards precision medicine initiatives, incorporating fetal sex and *O*-GlcNAcylation into clinical consideration can improve future diagnostic and targeted treatment methods for pregnancy complications.

## MATERIALS AND METHODS

### Placenta collection

Placenta samples were obtained from the Medical College of Wisconsin Maternal Research and Placenta and Cord Blood Bank (MCW MRPCB) under IRB approval. Gestational diabetes was diagnosed by the 100g oral glucose tolerance test following the Carpenter-Coustan criteria or if the 1-hour 50g glucose challenge test is > 200mg/dl. Placental samples were selected for this study on the following criteria (1) GDM diagnosis; (2) placed on insulin or oral hypoglycemic medication during pregnancy (3) no maternal history of hypertension (4) single gestation with known fetal sex without congenital anomalies. Placenta samples as the control group were comprised of women who tested negative for the GDM screening test with the 1-hour 50g glucose challenge test, but otherwise, the same exclusion criteria were applied.

### Cell culture

Bewo cells were maintained in Dulbecco’s modified Eagle’s medium (DMEM) containing 4.5 g/L (Standard, Std), 10% (v/v) fetal bovine serum (FBS), and 1% penicillin/streptomycin at 37°C in a 5% (v/v) CO_2_-enriched humidified atmosphere.

### *OGT* siRNA

MISSION siRNA targeting Human *OGT* were ordered at sigma (#EHU082301). Silencer™ Negative Control No. 1 siRNA (cat# AM4611 Thermofisher Scientific) was used as a control. Cells were transfected with the Lipofectamine RNAiMax (Thermo fisher scientific) according to the manufacturer protocol.

### RNA extraction

mRNA was isolated with the PureLink RNA Mini Kit (#12183018A, Thermo Fisher Scientific) following the manufacturer’s instruction. RNA integrity was verified by visual inspection of ribosomal RNA on agarose gels. RNA concentrations were measured with the LVis microplate on a FLUOstar Omega plate reader (BMG Labtech).

### cDNA preparation and qPCR

cDNA was then synthesized with SuperScript™ IV VILO™ Master Mix with ezDNase (#11766050, Thermofisher Scientific) according to the manufacturer’s instructions. qPCR reaction was then performed using PowerSYBR qPCR Master Mix (Thermo Fisher Scientific). Specific primers for each reaction are as follows: *ACTIN*-F: GATTCCTATGTGGGCGACGA; *ACTIN*-R: AGGTCTCAAACATGATCTGGGT; *OGT*-F: GGTGACTATGCCAGGAGAGACTCTTGC; *OGT*-R: CGAACTTTCTTCAGGTATTCTAGATC; *OGA-*F: CGGTGTGGTGGAAGGATTTTA; *OGA-*R: GTTGCTCAGCTTCCTCCACTG; *HCG*-F: GCTACTGCCCCACCATGACC; *HCG*-R: ATGGACTCGAAGCGCACATC; *GH2*-F: AAACGCTGATGTGGAGGCTG; *GH2*-R: GCCCGTAGTTCTTGAGCAGT; *LEP*-F: CTGTGCGGATTCTTGTGGCT; *LEP*-R: GAGGAGACTGACTGCGTGTGT; PL-F: TGACACCTACCAGGAGTTTGAAG; *PL*-R: GGGGTCACAGGATGCTACTC. qPCR was run on a QuantStudio 3 instrument (Applied Biosystems) with the recommended settings. The data were collected and processed on DataConnect (Thermo Fisher Scientific), and 2^−DDCq^ were calculated and plotted using Prism 9 (GraphPad Software).

### Protein lysis, SDS-PAGE, and Western Blotting

Samples were lysed in RIPA lysis buffer [10mM Tris-HCl, 150mM NaCl, 1% Triton X-100 (v/v), 0.5% NaDOC (w/v), 0.1% SDS (w/v), and protease inhibitors; pH7.5], vortexed and centrifuged at 18,000 g for 10 minutes at 4C. Sample lysates were resolved on 8% tris-glycine and transferred onto nitrocellulose. Membranes were then washed with ultra-purified water and labeled with No-Stain Protein Labeling Reagent (Thermo Fisher Scientific) according to kit instructions. Next, the membranes were blocked for 45 minutes with 5% (w/v) non-fat milk in Tris-buffered saline-Tween 20 buffer (TBS-T). Primary antibodies were added to the blocking solution, and the blots were incubated overnight at 4C with gentle agitation. Following primary incubation, blots were washed three times with 10mL of TBS-T for 10 minutes and incubated with anti-mouse and anti-rabbit fluorescent-conjugated secondary antibodies in a 1:20,000 dilution for 1 hour at room temperature. Three additional TBS-T washes with 10mL in 10 minutes were performed, and the blot signal was captured using Odyssey Fc (LI-COR).

### Antibodies

Antibodies for western blotting are used as follow: anti-*O*-GlcNAc (#ab2739-Abcam): 1:1000; Anti-actin (#A2066, Sigma Aldrich), 1:1000.

### RNA sequencing

Total RNA samples were quantified using Qubit 2.0 Fluorometer (Life Technologies, Carlsbad, CA, USA), and RNA integrity was checked using Agilent TapeStation 4200 (Agilent Technologies, Palo Alto, CA, USA). RNA sequencing libraries were prepared using the NEBNext Ultra RNA Library Prep Kit for Illumina, following the manufacturer’s instructions (NEB, Ipswich, MA, USA). Briefly, mRNAs were first enriched with Oligo(dT) beads. Enriched mRNAs were fragmented for 15 minutes at 94 °C. First-strand and second-strand cDNAs were subsequently synthesized. cDNA fragments were end-repaired and adenylated at 3’ends, and universal adapters were ligated to cDNA fragments, followed by index addition and library enrichment by limited-cycle PCR. The sequencing libraries were validated on the Agilent TapeStation (Agilent Technologies, Palo Alto, CA, USA) and quantified by using Qubit 2.0 Fluorometer (Invitrogen, Carlsbad, CA) as well as by quantitative PCR (KAPA Biosystems, Wilmington, MA, USA). Samples were processed on an Illumina HiSeq platform by GENEWIZ (Azenta Life Science).

Sequence reads were trimmed to remove possible adapter sequences and nucleotides with poor quality using Trimmomatic v.0.36. The trimmed reads were mapped to the Homo sapiens GRCh38 reference genome available on ENSEMBL using the STAR aligner v.2.5.2b. The STAR aligner is a splice aligner that detects splice junctions and incorporates them to help align the entire read sequences. BAM files were generated as a result of this step. Statistics of mapping the reads to the reference genome can be found in Table S1. Unique gene hit counts were calculated by using featureCounts from the Subread package v.1.5.2. Only unique reads that fell within exon regions were counted. After extraction of gene hit counts, the gene hit counts table was used for downstream differential expression analysis. Using DESeq2, a comparison of gene expression between the various groups was performed. The Wald test was used to generate p-values and log2 fold changes. Genes with an adjusted p-value < 0.05 were called differentially expressed genes for each comparison.

### GO enrichment analysis

BinGO app within Cytoscape was used to perform GO enrichment analysis. The goa_human GO list was used to cluster the set of genes based on their biological processes and determine their statistical significance.

### Data availability

RNA-sequencing raw (*R1.fastq and *R2.fastq) and processed (*.bam and counts.txt) files were deposited in Gene Expression Omnibus (GEO) repository under GSE200983.

## STATISTICS

Statistical analyses were performed on prism and R. Error bars always represent mean ± SEM. Various two-tails tests were used across the manuscript. Usually, for clinical sample a Mann– Whitney U test was performed to measure significance (unpaired, non-parametric) ^111^ and Pearson R was computed for correlation analysis. For RNA-Sequencing, Log2FoldChange: The Log2 fold change was calculated as follow: Log2(Group 2 mean normalized counts/Group 1 mean normalized counts). The Wald test was used to generate p-values and the Benjamini-Hochberg correction to obtain the adjusted p-value ^112^. For all other, classic student t-test (unpaired, parametric) was performed. Statistical significance was always represented as follow: ns ≥ 0.05, * p < 0.05, ** p < 0.01, *** p < 0.001, **** p < 0.0001.

## STUDY APPROVAL

Placenta samples were obtained from the Medical College of Wisconsin Maternal Research and Placenta and Cord Blood Bank (MCW MRPCB) under IRB approval. Written informed consent was received prior to participation.

## AUTHOR CONTRIBUTION

YC and SOVS designed the experiments and YC performed them. MC and AP provided the maternal-fetal medicine clinical expertise. All authors contributed to the manuscript.

## ACKNOWLEDGMENTS

We thank john Corbett, PhD, Meredith Cruz, MD, and the other member of the OVS lab for their valuable feedback along this project. We also thank Jennifer McIntosh, DO and Mary Rau for their help with the MCW placental bank. Funding for this study has been provided by the Women’s Health Research Program at MCW and supported by the Translational Glyc*O*mics Program for Career Development in Glycoscience at Milwaukee, WI.

## FIGURE LEGENDS

**Figure S1:**
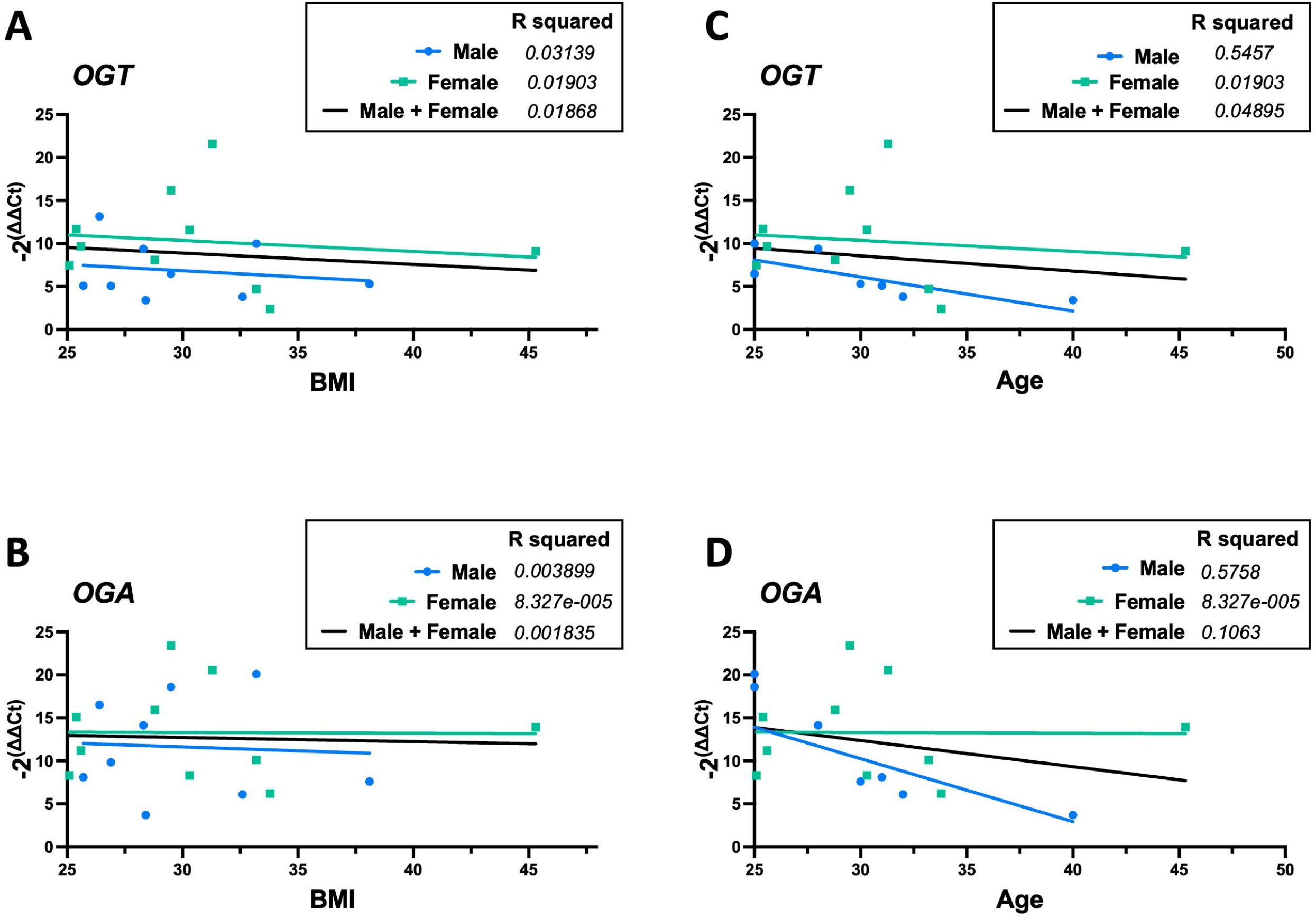
*OGT* and *OGA* expression do not correlate with BMI in male and female placentas. Pearson correlations were computed between BMI and *OGT* or *OGA* relative expression. ns p>0.05.

**Figure S2:**
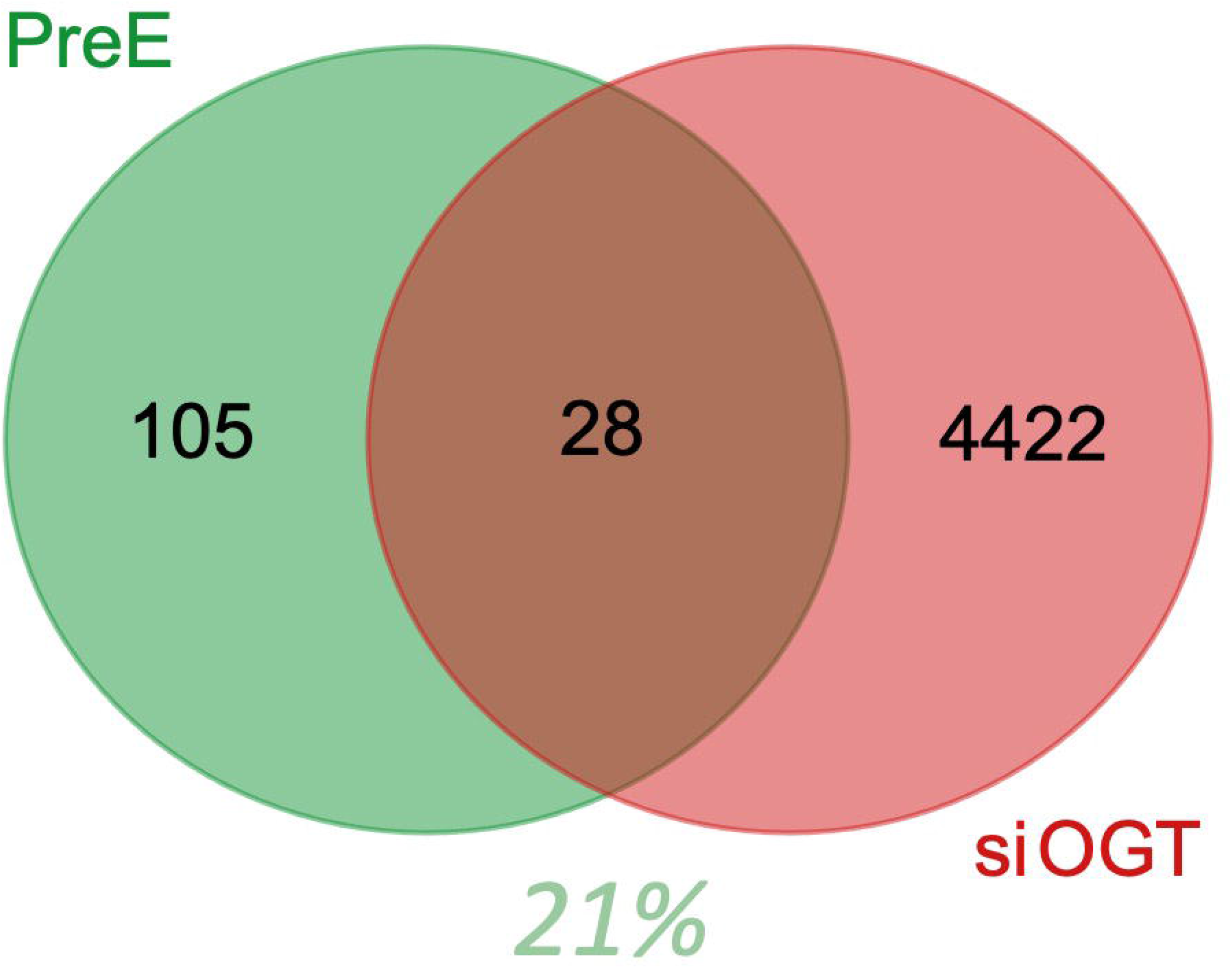
28 genes deregulated in si*OGT* overlap with preeclampsia. Preeclampsia genes were found in the Alliance of Genome Resources under DOID: 10591.

**Figure S3:**
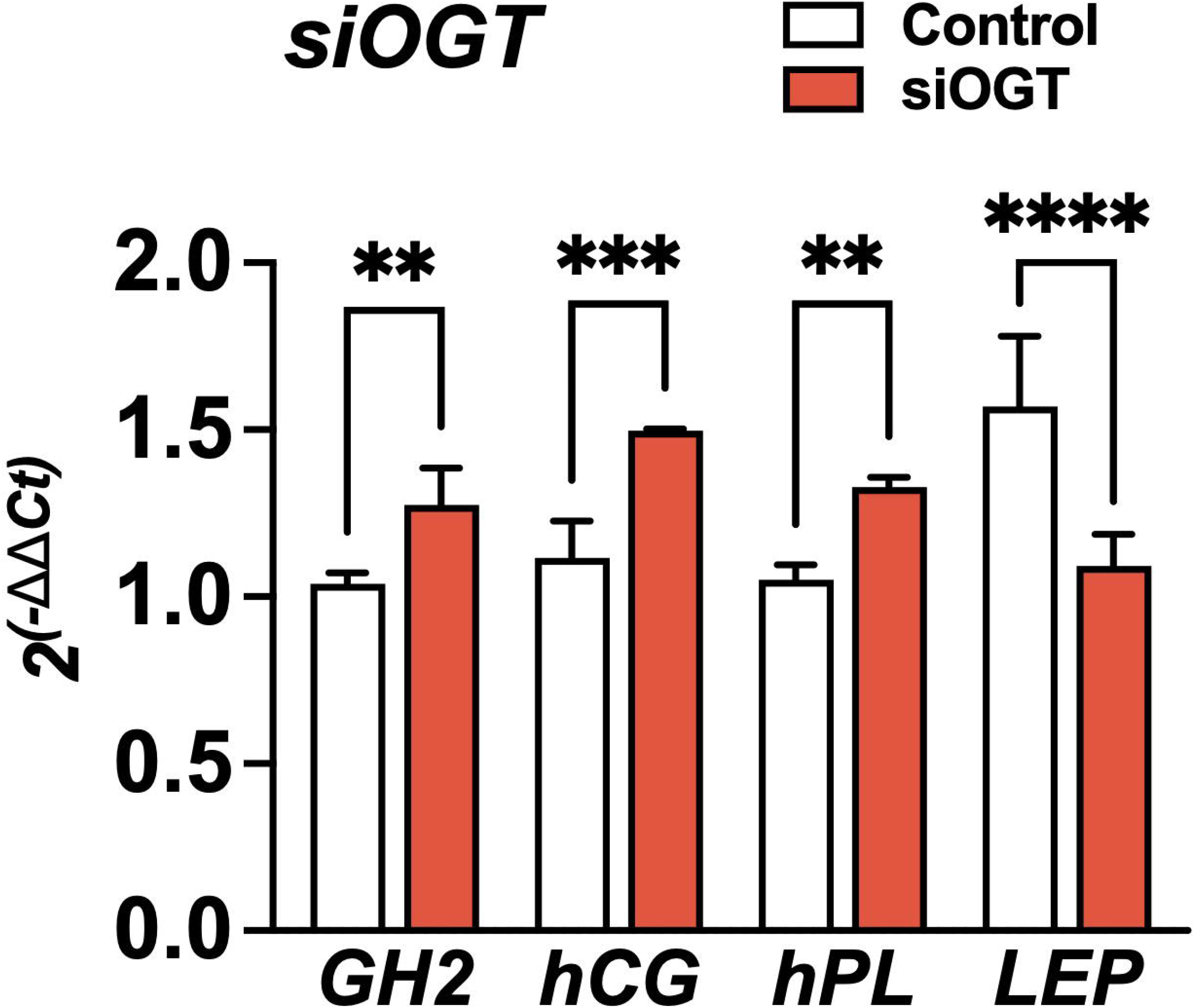
siOGT impacts hormonal levels in BeWo cells. *GH2, hCG, hPL*, and *LEP* were measured by qPCR after 48h of si*OGT* or siCtrl. Relative expression was calculated using the -2ΔΔCt method. Significance was calculated by student t-test; ** p < 0.01, *** p < 0.001, **** p < 0.0001.

**<u>Table S1:</u> Quality control of RNA sequencing**.

**<u>Table S2:</u> Differentially expressed genes in siOGT vs. SiCtrl**.

**<u>Table S3:</u> GO enrichment analysis**.

**<u>Table S4:</u> Genes commonly deregulated with other GDM studies**.

## REFERENCES

1. Saravanan P, Diabetes in Pregnancy Working Group, Maternal Medicine Clinical Study Group, Royal College of Obstetricians and Gynaecologists, UK. Gestational diabetes: opportunities for improving maternal and child health. Lancet Diabetes Endocrinol. 2020;8(9):793–800. doi:10.1016/S2213-8587(20)30161-3

2. Gunderson EP, Chiang V, Pletcher MJ, et al. History of gestational diabetes mellitus and future risk of atherosclerosis in mid-life: the Coronary Artery Risk Development in Young Adults study. J Am Heart Assoc. 2014;3(2):e000490. doi:10.1161/JAHA.113.000490

3. Xu Y, Shen S, Sun L, Yang H, Jin B, Cao X. Metabolic syndrome risk after gestational diabetes: a systematic review and meta-analysis. PloS One. 2014;9(1):e87863. doi:10.1371/journal.pone.0087863

4. Shah BR, Retnakaran R, Booth GL. Increased risk of cardiovascular disease in young women following gestational diabetes mellitus. Diabetes Care. 2008;31(8):1668–1669. doi:10.2337/dc08-0706

5. Bentley-Lewis R. Late cardiovascular consequences of gestational diabetes mellitus. Semin Reprod Med. 2009;27(4):322–329. doi:10.1055/s-0029-1225260

6. Di Cianni G, Lacaria E, Lencioni C, Resi V. Preventing type 2 diabetes and cardiovascular disease in women with gestational diabetes - The evidence and potential strategies. Diabetes Res Clin Pract. 2018;145:184–192. doi:10.1016/j.diabres.2018.04.021

7. Billionnet C, Mitanchez D, Weill A, et al. Gestational diabetes and adverse perinatal outcomes from 716,152 births in France in 2012. Diabetologia. 2017;60(4):636–644. doi:10.1007/s00125-017-4206-6

8. Burlina S, Dalfrà MG, Lapolla A. Short- and long-term consequences for offspring exposed to maternal diabetes: a review. J Matern-Fetal Neonatal Med Off J Eur Assoc Perinat Med Fed Asia Ocean Perinat Soc Int Soc Perinat Obstet. 2019;32(4):687–694. doi:10.1080/14767058.2017.1387893

9. Lowe WL, Scholtens DM, Lowe LP, et al. Association of Gestational Diabetes With Maternal Disorders of Glucose Metabolism and Childhood Adiposity. JAMA. 2018;320(10):1005–1016. doi:10.1001/jama.2018.11628

10. Bianco ME, Josefson JL. Hyperglycemia During Pregnancy and Long-Term Offspring Outcomes. Curr Diab Rep. 2019;19(12):143. doi:10.1007/s11892-019-1267-6

11. Tam WH, Ma RCW, Ozaki R, et al. In Utero Exposure to Maternal Hyperglycemia Increases Childhood Cardiometabolic Risk in Offspring. Diabetes Care. 2017;40(5):679–686. doi:10.2337/dc16-2397

12. Retnakaran R, Shah BR. Fetal Sex and the Natural History of Maternal Risk of Diabetes During and After Pregnancy. J Clin Endocrinol Metab. 2015;100(7):2574–2580. doi:10.1210/jc.2015-1763

13. Ehrlich SF, Eskenazi B, Hedderson MM, Ferrara A. Sex ratio variations among the offspring of women with diabetes in pregnancy. Diabet Med J Br Diabet Assoc. 2012;29(9):e273–278. doi:10.1111/j.1464-5491.2012.03663.x

14. Wulff-Fuentes E, Berendt RR, Massman L, et al. The human O -GlcNAcome database and meta-analysis. Sci Data. 2021;8(1):25. doi:10.1038/s41597-021-00810-4

15. Harwood KR, Hanover JA. Nutrient-driven O-GlcNAc cycling -think globally but act locally. J Cell Sci. 2014;127(Pt 9):1857-1867. doi:10.1242/jcs.113233

16. Chang YH, Weng CL, Lin KI. O-GlcNAcylation and its role in the immune system. J Biomed Sci. 2020;27(1):57. doi:10.1186/s12929-020-00648-9

17. Hart GW. Three Decades of Research on O-GlcNAcylation - A Major Nutrient Sensor That Regulates Signaling, Transcription and Cellular Metabolism. Front Endocrinol. 2014;5:183. doi:10.3389/fendo.2014.00183

18. Pravata VM, Muha V, Gundogdu M, et al. Catalytic deficiency of O-GlcNAc transferase leads to X-linked intellectual disability. Proc Natl Acad Sci U S A. 2019;116(30):14961–14970. doi:10.1073/pnas.1900065116

19. Pravata VM, Gundogdu M, Bartual SG, et al. A missense mutation in the catalytic domain of O-GlcNAc transferase links perturbations in protein O-GlcNAcylation to X-linked intellectual disability. FEBS Lett. 2020;594(4):717–727. doi:10.1002/1873-3468.13640

20. Selvan N, George S, Serajee FJ, et al. O-GlcNAc transferase missense mutations linked to X-linked intellectual disability deregulate genes involved in cell fate determination and signaling. J Biol Chem. 2018;293(27):10810–10824. doi:10.1074/jbc.RA118.002583

21. Vaidyanathan K, Niranjan T, Selvan N, et al. Identification and characterization of a missense mutation in the O-linked β-N-acetylglucosamine (O-GlcNAc) transferase gene that segregates with X-linked intellectual disability. J Biol Chem. 2017;292(21):8948–8963. doi:10.1074/jbc.M116.771030

22. Willems AP, Gundogdu M, Kempers MJE, et al. Mutations in N-acetylglucosamine (O-GlcNAc) transferase in patients with X-linked intellectual disability. J Biol Chem. 2017;292(30):12621–12631. doi:10.1074/jbc.M117.790097

23. Olivier-Van Stichelen S, Hanover JA. X-inactivation normalizes O-GlcNAc transferase levels and generates an O-GlcNAc-depleted Barr body. Front Genet. 2014;5:256. doi:10.3389/fgene.2014.00256

24. Olivier-Van Stichelen S, Abramowitz LK, Hanover JA. X marks the spot: does it matter that O-GlcNAc transferase is an X-linked gene? Biochem Biophys Res Commun. 2014;453(2):201–207. doi:10.1016/j.bbrc.2014.06.068

25. Dubois A, Deuve JL, Navarro P, et al. Spontaneous reactivation of clusters of X-linked genes is associated with the plasticity of X-inactivation in mouse trophoblast stem cells. Stem Cells Dayt Ohio. 2014;32(2):377–390. doi:10.1002/stem.1557

26. Mugford JW, Starmer J, Williams RL Jr, et al. Evidence for Local Regulatory Control of Escape from Imprinted X Chromosome Inactivation. Genetics. Published online March 19, 2014. doi:10.1534/genetics.114.162800

27. Calabrese JM, Sun W, Song L, et al. Site-specific silencing of regulatory elements as a mechanism of X inactivation. Cell. 2012;151(5):951–963. doi:10.1016/j.cell.2012.10.037

28. Splinter E, de Wit E, Nora EP, et al. The inactive X chromosome adopts a unique three-dimensional conformation that is dependent on Xist RNA. Genes Dev. 2011;25(13):1371–1383. doi:10.1101/gad.633311

29. Howerton CL, Morgan CP, Fischer DB, Bale TL. O-GlcNAc transferase (OGT) as a placental biomarker of maternal stress and reprogramming of CNS gene transcription in development. Proc Natl Acad Sci U S A. 2013;110(13):5169–5174. doi:10.1073/pnas.1300065110

30. Liu W, Wang H, Xue X, et al. OGT-related mitochondrial motility is associated with sex differences and exercise effects in depression induced by prenatal exposure to glucocorticoids. J Affect Disord. 2018;226:203–215. doi:10.1016/j.jad.2017.09.053

31. Shafi R, Iyer SP, Ellies LG, et al. The O-GlcNAc transferase gene resides on the X chromosome and is essential for embryonic stem cell viability and mouse ontogeny. Proc Natl Acad Sci U S A. 2000;97(11):5735–5739. doi:10.1073/pnas.100471497

32. Mohan R, Jo S, Da Sol Chung E, et al. Pancreatic β-Cell O-GlcNAc Transferase Overexpression Increases Susceptibility to Metabolic Stressors in Female Mice. Cells. 2021;10(10):2801. doi:10.3390/cells10102801

33. Shi JJ, Liu HF, Hu T, et al. Danggui-Shaoyao-San improves cognitive impairment through inhibiting O-GlcNAc-modification of estrogen α receptor in female db/db mice. J Ethnopharmacol. 2021;281:114562. doi:10.1016/j.jep.2021.114562

34. Pantaleon M, Steane SE, McMahon K, Cuffe JSM, Moritz KM. Placental O-GlcNAc-transferase expression and interactions with the glucocorticoid receptor are sex specific and regulated by maternal corticosterone exposure in mice. Sci Rep. 2017;7(1):2017. doi:10.1038/s41598-017-01666-8

35. Sinclair DAR, Syrzycka M, Macauley MS, et al. Drosophila O-GlcNAc transferase (OGT) is encoded by the Polycomb group (PcG) gene, super sex combs (sxc). Proc Natl Acad Sci U S A. 2009;106(32):13427–13432. doi:10.1073/pnas.0904638106

36. Moore M, Avula N, Jo S, Beetch M, Alejandro EU. Disruption of O-Linked N-Acetylglucosamine Signaling in Placenta Induces Insulin Sensitivity in Female Offspring. Int J Mol Sci. 2021;22(13):6918. doi:10.3390/ijms22136918

37. Radaelli T, Varastehpour A, Catalano P, Hauguel-de Mouzon S. Gestational diabetes induces placental genes for chronic stress and inflammatory pathways. Diabetes. 2003;52(12):2951–2958. doi:10.2337/diabetes.52.12.2951

38. Zhao YH, Wang DP, Zhang LL, Zhang F, Wang DM, Zhang WY. Genomic expression profiles of blood and placenta reveal significant immune-related pathways and categories in Chinese women with gestational diabetes mellitus. Diabet Med. 2011;28(2):237–246. doi:10.1111/j.1464-5491.2010.03140.x

39. Enquobahrie DA, Williams MA, Qiu C, Meller M, Sorensen TK. Global placental gene expression in gestational diabetes mellitus. Am J Obstet Gynecol. 2009;200(2):206.e1-13. doi:10.1016/j.ajog.2008.08.022

40. Yang Y, Guo F, Peng Y, et al. Transcriptomic Profiling of Human Placenta in Gestational Diabetes Mellitus at the Single-Cell Level. Front Endocrinol. 2021;12:474. doi:10.3389/fendo.2021.679582

41. Barker DJ. The fetal and infant origins of adult disease. BMJ. 1990;301(6761):1111. doi:10.1136/bmj.301.6761.1111

42. Di Renzo GC, Rosati A, Sarti RD, Cruciani L, Cutuli AM. Does fetal sex affect pregnancy outcome? Gend Med. 2007;4(1):19–30. doi:10.1016/s1550-8579(07)80004-0

43. Vatten LJ, Skjaerven R. Offspring sex and pregnancy outcome by length of gestation. Early Hum Dev. 2004;76(1):47–54. doi:10.1016/j.earlhumdev.2003.10.006

44. Persson M, Fadl H. Perinatal outcome in relation to fetal sex in offspring to mothers with pre-gestational and gestational diabetes--a population-based study. Diabet Med J Br Diabet Assoc. 2014;31(9):1047–1054. doi:10.1111/dme.12479

45. OuYang H, Chen B, Abdulrahman AM, Li L, Wu N. Associations between Gestational Diabetes and Anxiety or Depression: A Systematic Review. J Diabetes Res. 2021;2021:9959779. doi:10.1155/2021/9959779

46. Feng Y, Feng Q, Qu H, et al. Stress adaptation is associated with insulin resistance in women with gestational diabetes mellitus. Nutr Diabetes. 2020;10(1):1–4. doi:10.1038/s41387-020-0107-8

47. Li H, Shen L, Song L, et al. Early age at menarche and gestational diabetes mellitus risk: Results from the Healthy Baby Cohort study. Diabetes Metab. 2017;43(3):248–252. doi:10.1016/j.diabet.2017.01.002

48. Zhang X, Zhao X, Huo L, et al. Risk prediction model of gestational diabetes mellitus based on nomogram in a Chinese population cohort study. Sci Rep. 2020;10(1):21223. doi:10.1038/s41598-020-78164-x

49. Retnakaran R, Kramer CK, Ye C, et al. Fetal sex and maternal risk of gestational diabetes mellitus: the impact of having a boy. Diabetes Care. 2015;38(5):844–851. doi:10.2337/dc14-2551

50. Howerton CL, Bale TL. Targeted placental deletion of OGT recapitulates the prenatal stress phenotype including hypothalamic mitochondrial dysfunction. Proc Natl Acad Sci U S A. 2014;111(26):9639–9644. doi:10.1073/pnas.1401203111

51. Malassiné A, Frendo J -L., Evain-Brion D. A comparison of placental development and endocrine functions between the human and mouse model. Hum Reprod Update. 2003;9(6):531–539. doi:10.1093/humupd/dmg043

52. Pasek RC, Gannon M. Advancements and challenges in generating accurate animal models of gestational diabetes mellitus. Am J Physiol -Endocrinol Metab. 2013;305(11):E1327–E1338. doi:10.1152/ajpendo.00425.2013

53. del Rincon JP, Iida K, Gaylinn BD, et al. Growth hormone regulation of p85alpha expression and phosphoinositide 3-kinase activity in adipose tissue: mechanism for growth hormone-mediated insulin resistance. Diabetes. 2007;56(6):1638–1646. doi:10.2337/db06-0299

54. Dominici FP, Cifone D, Bartke A, Turyn D. Loss of sensitivity to insulin at early events of the insulin signaling pathway in the liver of growth hormone-transgenic mice. J Endocrinol. 1999;161(3):383–392. doi:10.1677/joe.0.1610383

55. Beck P, Daughaday WH. Human Placental Lactogen: Studies of Its Acute Metabolic Effects and Disposition in Normal Man*. J Clin Invest. 1967;46(1):103–110.

56. Ryan EA, Enns L. Role of gestational hormones in the induction of insulin resistance. J Clin Endocrinol Metab. 1988;67(2):341–347. doi:10.1210/jcem-67-2-341

57. Leturque A, Burnol AF, Ferre P, Girard J. Pregnancy-induced insulin resistance in the rat: assessment by glucose clamp technique. Am J Physiol-Endocrinol Metab. 1984;246(1):E25–E31. doi:10.1152/ajpendo.1984.246.1.E25

58. Ryan EA, O’Sullivan MJ, Skyler JS. Insulin action during pregnancy. Studies with the euglycemic clamp technique. Diabetes. 1985;34(4):380–389. doi:10.2337/diab.34.4.380

59. Kim JK. Hyperinsulinemic-euglycemic clamp to assess insulin sensitivity in vivo. Methods Mol Biol Clifton NJ. 2009;560:221–238. doi:10.1007/978-1-59745-448-3_15

60. Ahmed-Sorour H, Bailey CJ. Role of ovarian hormones in the long-term control of glucose homeostasis. Interaction with insulin, glucagon and epinephrine. Horm Res. 1980;13(6):396–403. doi:10.1159/000179307

61. Takeda K, Toda K, Saibara T, et al. Progressive development of insulin resistance phenotype in male mice with complete aromatase (CYP19) deficiency. J Endocrinol. 2003;176(2):237–246. doi:10.1677/joe.0.1760237

62. Bryzgalova G, Gao H, Ahren B, et al. Evidence that oestrogen receptor-alpha plays an important role in the regulation of glucose homeostasis in mice: insulin sensitivity in the liver. Diabetologia. 2006;49(3):588–597. doi:10.1007/s00125-005-0105-3

63. Scott LJ, Bonnycastle LL, Willer CJ, et al. Association of transcription factor 7-like 2 (TCF7L2) variants with type 2 diabetes in a Finnish sample. Diabetes. 2006;55(9):2649–2653. doi:10.2337/db06-0341

64. El-Lebedy D, Ashmawy I. Common variants in TCF7L2 and CDKAL1 genes and risk of type 2 diabetes mellitus in Egyptians. J Genet Eng Biotechnol. 2016;14(2):247–251. doi:10.1016/j.jgeb.2016.10.004

65. Kirwan JP, Varastehpour A, Jing M, et al. Reversal of insulin resistance postpartum is linked to enhanced skeletal muscle insulin signaling. J Clin Endocrinol Metab. 2004;89(9):4678–4684. doi:10.1210/jc.2004-0749

66. Barbour LA, Shao J, Qiao L, et al. Human placental growth hormone increases expression of the p85 regulatory unit of phosphatidylinositol 3-kinase and triggers severe insulin resistance in skeletal muscle. Endocrinology. 2004;145(3):1144–1150. doi:10.1210/en.2003-1297

67. Bailey CJ, Ahmed-Sorour H. Role of ovarian hormones in the long-term control of glucose homeostasis. Diabetologia. 1980;19:475–481. doi:10.1007/BF00281829

68. Costrini NV, Kalkhoff RK. Relative effects of pregnancy, estradiol, and progesterone on plasma insulin and pancreatic islet insulin secretion. J Clin Invest. 1971;50(5):992–999.

69. Brelje TC, Scharp DW, Lacy PE, et al. Effect of homologous placental lactogens, prolactins, and growth hormones on islet B-cell division and insulin secretion in rat, mouse, and human islets: implication for placental lactogen regulation of islet function during pregnancy. Endocrinology. 1993;132(2):879–887. doi:10.1210/endo.132.2.8425500

70. Huang C, Snider F, Cross JC. Prolactin receptor is required for normal glucose homeostasis and modulation of beta-cell mass during pregnancy. Endocrinology. 2009;150(4):1618–1626. doi:10.1210/en.2008-1003

71. Fujinaka Y, Sipula D, Garcia-Ocaña A, Vasavada RC. Characterization of mice doubly transgenic for parathyroid hormone-related protein and murine placental lactogen: a novel role for placental lactogen in pancreatic beta-cell survival. Diabetes. 2004;53(12):3120–3130. doi:10.2337/diabetes.53.12.3120

72. N B, Jh N. The stimulatory effect of growth hormone, prolactin, and placental lactogen on beta-cell proliferation is not mediated by insulin-like growth factor-I. Endocrinology. 1991;129(2):883–888. doi:10.1210/endo-129-2-883

73. Nielsen JH. Effects of Growth Hormone, Prolactin, and Placental Lactogen on Insulin Content and Release, and Deoxyribonucleic Acid Synthesis in Cultured Pancreatic Islets. Endocrinology. 1982;110(2):600–606. doi:10.1210/endo-110-2-600

74. Brelje TC, Allaire P, Hegre O, Sorenson RL. Effect of prolactin versus growth hormone on islet function and the importance of using homologous mammosomatotropic hormones. Endocrinology. 1989;125(5):2392–2399. doi:10.1210/endo-125-5-2392

75. Weinhaus AJ, Stout LE, Sorenson RL. Glucokinase, hexokinase, glucose transporter 2, and glucose metabolism in islets during pregnancy and prolactin-treated islets in vitro: mechanisms for long term up-regulation of islets. Endocrinology. 1996;137(5):1640–1649. doi:10.1210/endo.137.5.8612496

76. Sorenson RL, Brelje TC. Adaptation of Islets of Langerhans to Pregnancy: β-Cell Growth, Enhanced Insulin Secretion and the Role of Lactogenic Hormones. Horm Metab Res. 1997;29(6):301–307. doi:10.1055/s-2007-979040

77. Arumugam R, Fleenor D, Freemark M. Knockdown of prolactin receptors in a pancreatic beta cell line: effects on DNA synthesis, apoptosis, and gene expression. Endocrine. 2014;46(3):568–576. doi:10.1007/s12020-013-0073-1

78. Dominici FP, Argentino DP, Muñoz MC, Miquet JG, Sotelo AI, Turyn D. Influence of the crosstalk between growth hormone and insulin signalling on the modulation of insulin sensitivity. Growth Horm IGF Res Off J Growth Horm Res Soc Int IGF Res Soc. 2005;15(5):324–336. doi:10.1016/j.ghir.2005.07.001

79. Napso T, Yong HEJ, Lopez-Tello J, Sferruzzi-Perri AN. The Role of Placental Hormones in Mediating Maternal Adaptations to Support Pregnancy and Lactation. Front Physiol. 2018;9:1091. doi:10.3389/fphys.2018.01091

80. Brănişteanu DD, Mathieu C. Progesterone in gestational diabetes mellitus: guilty or not guilty? Trends Endocrinol Metab TEM. 2003;14(2):54–56. doi:10.1016/s1043-2760(03)00003-1

81. Qi X, Gong B, Yu J, et al. Decreased cord blood estradiol levels in related to mothers with gestational diabetes. Medicine (Baltimore). 2017;96(21):e6962. doi:10.1097/MD.0000000000006962

82. Picard F, Wanatabe M, Schoonjans K, Lydon J, O’Malley BW, Auwerx J. Progesterone receptor knockout mice have an improved glucose homeostasis secondary to β-cell proliferation. Proc Natl Acad Sci. 2002;99(24):15644–15648. doi:10.1073/pnas.202612199

83. Li HP, Chen X, Li MQ. Gestational diabetes induces chronic hypoxia stress and excessive inflammatory response in murine placenta. Int J Clin Exp Pathol. 2013;6(4):650–659.

84. Rygaard K, Revol A, Esquivel-Escobedo D, Beck BL, Barrera-Saldaña HA. Absence of human placental lactogen and placental growth hormone (HGH-V) during pregnancy: PCR analysis of the deletion. Hum Genet. 1998;102(1):87–92. doi:10.1007/s004390050658

85. Le TN, Elsea SH, Romero R, Chaiworapongsa T, Francis GL. Prolactin receptor gene polymorphisms are associated with gestational diabetes. Genet Test Mol Biomark. 2013;17(7):567–571. doi:10.1089/gtmb.2013.0009

86. Ekinci EI, Torkamani N, Ramchand SK, et al. Higher maternal serum prolactin levels are associated with reduced glucose tolerance during pregnancy. J Diabetes Investig. 2017;8(5):697–700. doi:10.1111/jdi.12634

87. Buschur E, Stetson B, Barbour LA. Diabetes In Pregnancy. In: Feingold KR, Anawalt B, Boyce A, et al., eds. Endotext. MDText.com, Inc.; 2000. Accessed May 9, 2020. http://www.ncbi.nlm.nih.gov/books/NBK279010/

88. Handwerger S, Freemark M. The roles of placental growth hormone and placental lactogen in the regulation of human fetal growth and development. J Pediatr Endocrinol Metab JPEM. 2000;13(4):343–356. doi:10.1515/jpem.2000.13.4.343

89. Arumugam R, Horowitz E, Lu D, et al. The interplay of prolactin and the glucocorticoids in the regulation of beta-cell gene expression, fatty acid oxidation, and glucose-stimulated insulin secretion: implications for carbohydrate metabolism in pregnancy. Endocrinology. 2008;149(11):5401–5414. doi:10.1210/en.2008-0051

90. Fujinaka Y, Takane K, Yamashita H, Vasavada RC. Lactogens promote beta cell survival through JAK2/STAT5 activation and Bcl-XL upregulation. J Biol Chem. 2007;282(42):30707–30717. doi:10.1074/jbc.M702607200

91. Guerre-Millo M. Extending the glucose/fatty acid cycle: a glucose/adipose tissue cycle. Biochem Soc Trans. 2003;31(Pt 6):1161-1164. doi:10.1042/bst0311161

92. Perry RJ, Zhang XM, Zhang D, et al. Leptin reverses diabetes by suppression of the hypothalamic-pituitary-adrenal axis. Nat Med. 2014;20(7):759–763. doi:10.1038/nm.3579

93. Masuzaki H, Ogawa Y, Sagawa N, et al. Nonadipose tissue production of leptin: leptin as a novel placenta-derived hormone in humans. Nat Med. 1997;3(9):1029–1033. doi:10.1038/nm0997-1029

94. Yang Y, Wu N. Gestational Diabetes Mellitus and Preeclampsia: Correlation and Influencing Factors. Front Cardiovasc Med. 2022;9. Accessed May 6, 2022. https://www.frontiersin.org/article/10.3389/fcvm.2022.831297

95. Liu J, Shao X, Qin W, et al. Quantitative chemoproteomics reveals O-GlcNAcylation of cystathionine γ-lyase (CSE) represses trophoblast syncytialization. Cell Chem Biol. 2021;28(6):788-801.e5. doi:10.1016/j.chembiol.2021.01.024

96. Llamas-Covarrubias MA, Valle Y, Navarro-Hernández RE, et al. Serum levels of macrophage migration inhibitory factor are associated with rheumatoid arthritis course. Rheumatol Int. 2012;32(8):2307–2311. doi:10.1007/s00296-011-1951-6

97. Kleemann R, Bucala R. Macrophage migration inhibitory factor: critical role in obesity, insulin resistance, and associated comorbidities. Mediators Inflamm. 2010;2010:610479. doi:10.1155/2010/610479

98. Stojanovic I, Saksida T, Stosic-Grujicic S. Beta cell function: the role of macrophage migration inhibitory factor. Immunol Res. 2012;52(1-2):81–88. doi:10.1007/s12026-012-8281-y

99. Finucane OM, Reynolds CM, McGillicuddy FC, Roche HM. Insights into the role of macrophage migration inhibitory factor in obesity and insulin resistance. Proc Nutr Soc. 2012;71(4):622–633. doi:10.1017/S0029665112000730

100. Sánchez-Zamora YI, Rodriguez-Sosa M. The role of MIF in type 1 and type 2 diabetes mellitus. J Diabetes Res. 2014;2014:804519. doi:10.1155/2014/804519

101. Stojanovic I, Saksida T, Timotijevic G, Sandler S, Stosic-Grujicic S. Macrophage migration inhibitory factor (MIF) enhances palmitic acid- and glucose-induced murine beta cell dysfunction and destruction in vitro. Growth Factors Chur Switz. 2012;30(6):385–393. doi:10.3109/08977194.2012.734506

102. Verschuren L, Kooistra T, Bernhagen J, et al. MIF deficiency reduces chronic inflammation in white adipose tissue and impairs the development of insulin resistance, glucose intolerance, and associated atherosclerotic disease. Circ Res. 2009;105(1):99–107. doi:10.1161/CIRCRESAHA.109.199166

103. Stojanovic I, Saksida T, Nikolic I, Nicoletti F, Stosic-Grujicic S. Macrophage migration inhibitory factor deficiency protects pancreatic islets from cytokine-induced apoptosis in vitro. Clin Exp Immunol. 2012;169(2):156–163. doi:10.1111/j.1365-2249.2012.04607.x

104. Solomon SS, Majumdar G, Martinez-Hernandez A, Raghow R. A critical role of Sp1 transcription factor in regulating gene expression in response to insulin and other hormones. Life Sci. 2008;83(9-10):305-312. doi:10.1016/j.lfs.2008.06.024

105. LiCalsi C, Christophe S, Steger DJ, Buescher M, Fischer W, Mellon PL. AP-2 family members regulate basal and cAMP-induced expression of human chorionic gonadotropin. Nucleic Acids Res. 2000;28(4):1036–1043. doi:10.1093/nar/28.4.1036

106. Pena P, Reutens AT, Albanese C, et al. Activator protein-2 mediates transcriptional activation of the CYP11A1 gene by interaction with Sp1 rather than binding to DNA. Mol Endocrinol Baltim Md. 1999;13(8):1402–1416. doi:10.1210/mend.13.8.0335

107. Piao YS, Peltoketo H, Vihko P, Vihko R. The proximal promoter region of the gene encoding human 17beta-hydroxysteroid dehydrogenase type 1 contains GATA, AP-2, and Sp1 response elements: analysis of promoter function in choriocarcinoma cells. Endocrinology. 1997;138(8):3417–3425. doi:10.1210/endo.138.8.5329

108. Sheridan RM, Stanek J, Khoury J, Handwerger S. Abnormal expression of transcription factor AP-2α in pathologic placentas. Hum Pathol. 2012;43(11):1866–1874. doi:10.1016/j.humpath.2012.01.011

109. Xie X, Wu Q, Zhang K, et al. O-GlcNAc modification regulates MTA1 transcriptional activity during breast cancer cell genotoxic adaptation. Biochim Biophys Acta Gen Subj. 2021;1865(8):129930. doi:10.1016/j.bbagen.2021.129930

110. Wulff-Fuentes E, Berendt RR, Massman L, et al. The human O-GlcNAcome database and meta-analysis. Sci Data. 2021;8(1):25. doi:10.1038/s41597-021-00810-4

111. Sundjaja JH, Shrestha R, Krishan K. McNemar And Mann-Whitney U Tests. In:StatPearls. StatPearls Publishing; 2022. Accessed May 11, 2022. http://www.ncbi.nlm.nih.gov/books/NBK560699/

112. Varet H, Brillet-Guéguen L, Coppée JY, Dillies MA. SARTools: A DESeq2- and EdgeR-Based R Pipeline for Comprehensive Differential Analysis of RNA-Seq Data. PloS One. 2016;11(6):e0157022. doi:10.1371/journal.pone.0157022

